# An RFX-driven ciliogenic cell state defines ex vivo FGFR-inhibitor sensitivity in colorectal cancer

**DOI:** 10.64898/2026.10.12.756637

**Authors:** Lucie Kralickova, Nicola Miglino, Cinzia Esposito, Ramon Pfaendler, Rebekka Wegmann, Pilar Ballesteros Cuartero, Julien Mena, Federica Panebianco, Laura A Boos, Ulrich Richter, Parisa Rahimzadeh, Tarun Mehra, Viktor Hendrik Koelzer, Salvatore Piscuoglio, Andreas Wicki, Berend Snijder

**Affiliations:** Institute of Molecular Systems Biology, Department of Biology, ETH Zurich, Zü rich, Switzerland; Botnar Institute of Immune Engineering, Basel, Switzerland; Department of Medical Oncology and Hematology, University of Zurich and University Hospital, Zurich, Switzerland; Department of Biomedicine, University of Basel, Basel, Switzerland; Computational and Translational Pathology (CTP) Lab, Department of Biomedical Engineering, University of Basel, Switzerland; Institute of Medical Genetics and Pathology, University Hospital Basel, Basel, Switzerland; IRCCS Humanitas Research Hospital, Milan, Italy; Swiss Institute of Bioinformatics, Lausanne, Switzerland; Comprehensive Cancer Center Zurich, Zü rich, Switzerland; Department of Biomedical Sciences, Humanitas University, Milan, Italy; Medical and Scientific Affairs, Thermo Fisher Scientific, Waltham, Massachusetts, USA

**Author notes:** Corresponding author eering. Shared last (Andreas Wicki, Berend Snijder).

## Abstract

Genomic stratification leaves most patients with colorectal cancer (CRC) without opportunities for targeted therapy. Seeking tumour state-linked drug sensitivities, we combined image-based ex vivo drug profiling with matched transcriptomics in 127 prospectively collected patient biopsies across CRC stages and molecular subtypes. Network-level integration identified a ciliogenic cell state defined by upregulation of cilia related genes with selective sensitivity to the FGFR inhibitor erdafitinib. This ciliogenic signature was associated with poor progression-free survival (hazard ratio 2.28; adjusted p=0.0098), even in early-stage colorectal cancer. Furthermore, knockdown of the transcriptional regulators RFX2 and RFX3, upstream of the ciliogenic cell state, attenuated erdafitinib sensitivity. Orthogonal bulk and single-cell analyses across normal, primary and metastatic tissues contextualised the ciliogenic programme along a transition from canonical epithelial states towards fetal and non-canonical tumour states, concomitant with Wnt-related remodelling, FGFR1/FGFR2 upregulation, and loss of intestinal epithelial identity. Together, these findings define a tumour-intrinsic ciliogenic cell state as a feature of CRC progression, revealing a patient-stratified repurposing opportunity for targeted therapy beyond the current CRC treatment paradigms.

## 1 Main

Precision oncology in colorectal cancer (CRC) remains anchored to a small set of genomic biomarkers, leaving most patients without an actionable targeted option [1, 2]. Yet tumour progression is driven not only by genetic alterations but also by dynamic, non-genetic shifts in cellular state [3, 4, 5, 6, 7]. In colorectal cancer, malignant cells lose canonical intestinal identity and adopt developmental or non-canonical transcriptional programmes, generating substantial intra- and inter-tumour heterogeneity [8, 9]. Such cell-state diversity is increasingly recognised as a key determinant of tumour behaviour, yet its contribution to therapeutic vulnerability remains incompletely understood [10, 11, 5]. To connect these cell states to actionable vulnerabilities, functional profiling of fresh patient tumour material offers a complementary route that directly connects tumour phenotype to drug response [12, 13, 14, 15]. Applying this across a prospective colorectal cancer cohort, we identify a ciliogenic transcriptional state selectively sensitive to the FGFR inhibitor erdafitinib, that extends FGFR-directed therapy to a colorectal cancer population beyond those defined by FGFR genomic alteration.

## 2 Results

### Functional profiling in CRC biopsies delineates drug response-associated transcriptional programmes

We established an image-based ex vivo drug-screening platform for the functional characterisation of patient-derived colorectal cancer (CRC) tissue, and profiled 149 samples from 148 patients including both surgical resections and core needle biopsies (Figure 1a; Supplementary Figure 1a). Drug-response profiles were successfully obtained for 137 samples (92%; Supplementary Figure 2a), demonstrating that robust functional profiling is feasible in freshly dissociated, clinically accessible CRC material,including core needle biopsies obtained as part of clinical routine. This provides a complementary functional readout alongside approaches based on long-term ex vivo patient-derived cultures [16, 17]. After applying prespecified cohort and sample inclusion criteria, 127 patient biopsies across disease stages and molecular subtypes were retained for downstream analysis (Supplementary Figure 2a-c).

**Figure 1:**
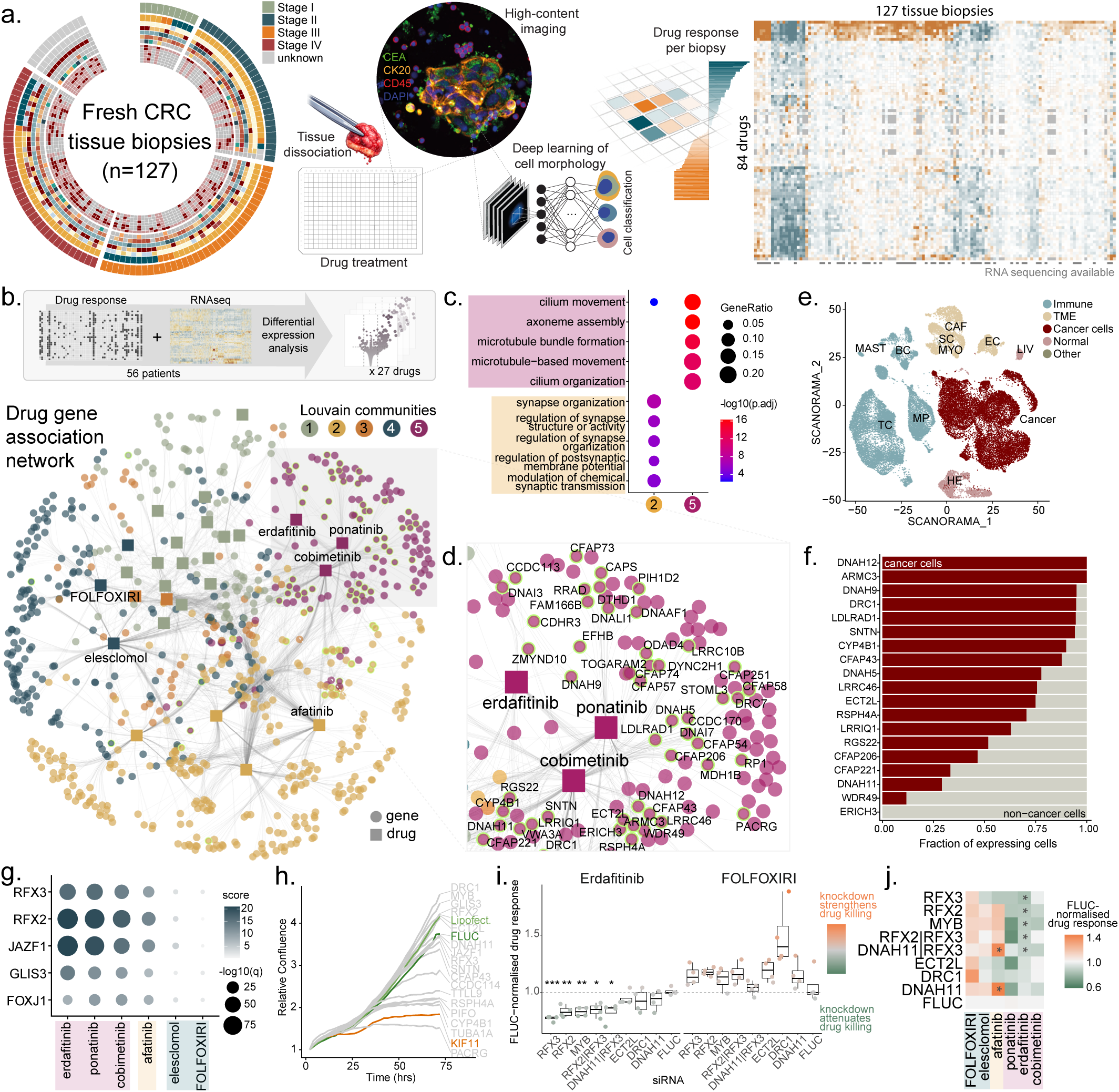
Functional profiling identifies a tumour-intrinsic ciliogenic cell state linked to selective erdafitinib sensitivity. **a.** Study overview. A cohort of 127 fresh colorectal cancer (CRC) biopsies spanning all clinical stages was profiled using an image-based ex vivo drug screening platform combined with deep-learning-based cell morphology classification, generating functional responses across 84 compounds and compound combinations. Matched RNA sequencing was generated for a subset of samples (n = 56). Detailed cohort characteristics and drug response metrics are provided in Supplementary. **b.** Gene-drug association network derived from DESeq2 differential expression analysis. Bipartite network integrating differential gene expression results across all drugs. Nodes represent drugs (squares) or genes (circles). Edges connect a drug to genes significantly upregulated in ex vivo drug responders versus non-responders based on a Wald test with Benjamini-Hochberg correction (shrunken log_2_ fold change *>* 1, adjusted *p <* 0.05), restricted to the top 300 genes per drug ranked by effect size. Node colour denotes Louvain community membership identified by maximisation of network modularity. Genes overlapping a curated 500-gene cilia reference set (CG500) are highlighted (green outline). **c.** Gene Ontology (GO) enrichment of Louvain communities shown in **b**. Biological Process (BP) terms were tested per community. Dot size indicates gene ratio. Colour indicates statistical significance (*−* log_10_ adjusted *p*-value). Only communities with significant enrichment (adjusted *p <* 0.05; *p*-value cut-off *<* 0.01) are shown. For each community, the top five GO terms ranked by adjusted *p*-value are displayed. **d.** Zoom-in of Louvain Community 5 from **b**, enriched for cilia-associated genes. **e.** UMAP with Scanorama integration showing cell identities of 34,895 cells from 26 CRC patients, as previously described [19]. Immune: TC, T cell-like; MP, antigen-presenting cells; BC, B cell-like; MAST, mast-like cells. Tumour microenvironment (TME): CAF, fibroblasts, myofibroblasts and pericytes; EC, endothelial cells. Cancer cells: CC, epithelial cancer cells. Normal: HE, healthy epithelial cells; LIV, liver cells (hepatocytes and cholangiocytes). Other: MYO, myoblasts/myocytes; SC, Schwann cells. **f.** Single-cell expression of cilia-associated genes derived from Louvain Community 5. The top 19 genes were selected based on mean shrunken log_2_ fold change across erdafitinib, cobimetinib and ponatinib responders. Bars indicate the fraction of expressing cells (normalised expression *>* 0), stratified by cancer versus non-cancer cells. **g.** Transcription factor activity inferred from differential expression analysis. Shown are transcription factors significantly enriched in common across cobimetinib, erdafitinib, and ponatinib (adjusted *p <* 0.01). Colour intensity reflects absolute activity score and bubble size indicates statistical significance (*−* log_10_ adjusted *p*-value). **h.** Time course of GP2D cell confluence following siRNA knockdown, measured by live-cell imaging (n = 3 technical replicates per siRNA). **i.** Drug response of GP2D cells to erdafitinib and FOLFOXIRI following siRNA knockdown, shown as FLUC-normalised viable (cleaved caspase-3–negative) cell number relative to DMSO. Values *<* 1 indicate attenuated drug response. Each point represents one technical replicate. **j.** Heatmap of the median FLUC-normalised drug response (as in **i**, n=4) across all tested conditions. Unless otherwise indicated, significance was assessed using a two-tailed Student’s *t*-test with Benjamini-Hochberg correction. Significance thresholds: *q <* 0.05 (*), *q <* 0.01 (**), *q <* 0.001 (***), *q <* 0.0001 (****).

To systematically quantify drug responses across this large image-based dataset, we applied deep-learning-assisted single-cell analyses to distinguish malignant from non-malignant cells (Supplementary Figure 3a-f) and measure cell-type-specific drug effects across 84 drugs or drug combinations (Supplementary Figure 4a). By focusing on malignant-cell-selective responses, this approach favours tumour-selective effects over collateral toxicity (Supplementary Figure 5a-b). This generated a patient-by-drug response matrix that captured substantial inter-patient heterogeneity and provided the basis for downstream integration of functional and transcriptomic data (Supplementary Figures 4a, 5c-f).

To identify transcriptional programmes associated with the single-cell resolved drug response profiles, we generated transcriptomic data from 56 matched biopsies and integrated this data with the functional profiles using drug-wise differential expression analysis (Figure 1b, Supplementary Figure 6a-d). Significantly associated gene-drug pairs were assembled into an association network and partitioned by Louvain clustering, revealing five communities (Figure 1b; Supplementary Figure 7a). Two of these showed clear functional enrichment: a synapse-related programme (community 2) associated with afatinib, osimertinib, dasatinib and crizotinib sensitivity, and a cilia-related programme (community 5) associated with erdafitinib, ponatinib and cobimetinib (Figure 1c-d). Among the identified communities, the cilia-enriched programme, surprisingly marked by motile cilia-related CFAP and DNAH family members [18], had the strongest enrichment signal and highest density of related genes, prioritising it as a candidate tumour cell state linked to selective therapeutic vulnerability ex vivo.

### A ciliogenic transcriptional cell state is functionally linked to selective erdafitinib sensitivity

To determine whether the cilia gene-enriched community reflected a tumour-intrinsic programme, we examined its cellular origin in CRC single-cell RNA-seq data [19]. Although the top drug-associated cilia genes from community 5, including DNAH12, DRC1, and CFAP43, were generally lowly expressed, they were preferentially detected in cancer cells rather than non-cancer cells, indicating that this signal arises predominantly from the tumour compartment (Figure 1e-f; Supplementary Figure 8a). Consistently, similar cilia-associated malignant cell states have recently been recognised in colorectal cancer and glioblastoma [8, 20].

To test the functional relevance of this cilia-enriched programme, we inferred transcription factor activity to nominate candidate upstream regulators. This identified increased activity of cilia-associated transcription factors [21] RFX2 and RFX3 in association with response to erdafitinib, ponatinib and cobimetinib (Figure 1g). Functional perturbation showed that several cilia-associated knockdowns reduced the fitness of colorectal cancer cells (Figure 1h, Supplementary Figure 9a-b). Most importantly, depletion of RFX2, RFX3 and MYB attenuated response to FGFR-inhibitor erdafitinib in this context, whereas response to the standard-of-care chemotherapy FOLFOXIRI, assigned to a different transcriptional community, remained unchanged (Figure 1i; Supplementary Figure 9c). Similar attenuation trends were observed for the multityrosine kinase inhibitors ponatinib and cobimetinib, but not for negative-control drugs from other communities, including elesclomol and afatinib (Figure 1j; Supplementary Figure 9c). Together, these data suggest that the ciliogenic programme, i.e., the transcriptional upregulation of cilia-related genes, reflects a tumour-intrinsic cell state functionally coupled to FGFR inhibitor sensitivity.

### The ciliogenic signature is clinically prognostic and progression-associated in CRC

Given the incomplete annotation of cilia-associated genes in classical gene ontology, we identified the gene cluster of more broadly cilia-annotated genes [21] that was most co-expressed with community 5, termed the cancer-associated ciliogenic signature (CCGS; Figure 2a). We then asked whether CCGS expression captured a clinically meaningful CRC subset beyond the discovery cohort. In The Cancer Genome Atlas (TCGA) CRC cohort (n=592), high CCGS expression was associated with poor progression-free survival (hazard ratio [HR] = 2.28, adjusted *p* = 0.0098; Figure 2c), whereas high expression of unselected cilia-related genes (CG500; HR = 0.75, adjusted *p* = 0.19; Figure 2d) as well as the cilia-related genes that are not part of CCGS (ciliaOTHER; HR = 0.69, adjusted *p* = 0.18; Figure 2e) rather associated with better prognosis. Thus, prognostic information resides in a specific cell-state-associated cilia gene subset rather than in generic ciliogenesis-associated expression. Although CCGS increased with pathological stage, stage-specific analyses showed that its prognostic value is apparent in earlier-stage tumours, arguing that this ciliogenic cell state is not simply a surrogate for advanced disease (Figure 2f-h; Supplementary Figure 10ab,d-g). Together, these findings establish CCGS as a refined transcriptional signature of a progression-associated CRC cell state that is clinically prognostic.

**Figure 2:**
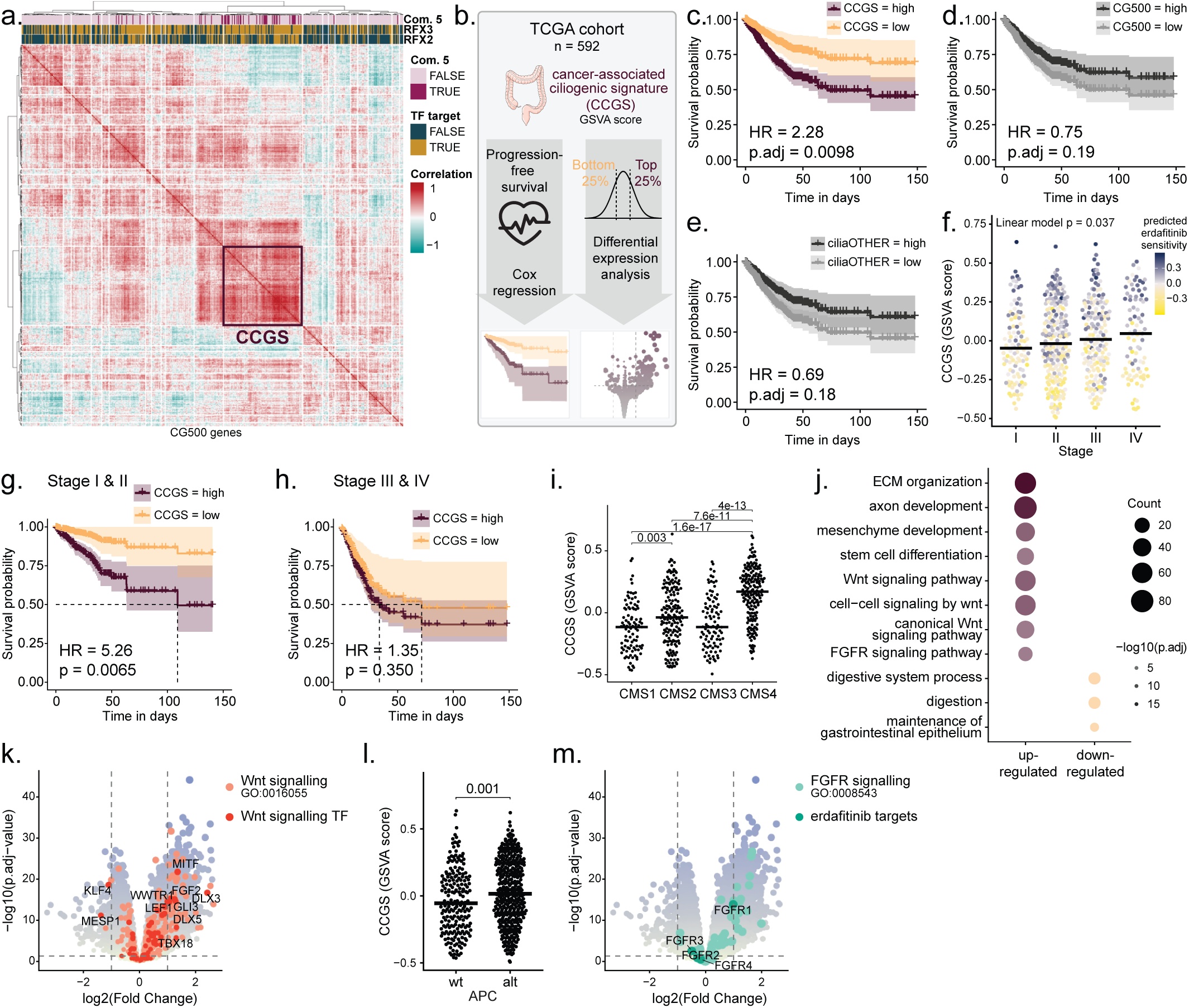
A refined transcriptional readout captures a prognostic ciliogenic cell state embedded in Wnt-associated CRC remodelling. **a.** Co-expression structure of curated cilia genes in the IBDS RNA-seq cohort (n = 56). Expression of the CG500 cilia reference set was used to compute a gene–gene Spearman correlation matrix across samples, followed by hierarchical clustering to identify co-regulated modules. Genes are annotated for overlap with the Louvain community 5 cilia-associated genes identified from drug-responder differential expression (Fig. 1b-d), and as RFX2 and RFX3 targets from CilioGenics [21]. The module exhibiting the greatest overlap with Louvain community 5 was selected as the refined cancer-associated ciliogenic signature (CCGS; n = 99 genes). **b.** Schematic overview of validation analyses in the TCGA colorectal cancer cohort (n = 592). Bulk RNA-seq data were used to compute GSVA scores for the CCGS and related cilia gene sets. These scores were evaluated for associations with progression-free survival (PFS) and used to contrast transcriptional profiles between high- and low-scoring tumours (top and bottom 25%, respectively). **c-e.** Kaplan–Meier (KM) plots of PFS in the TCGA cohort, stratified into high and low groups for CCGS (**c.**), CG500 (**d.**) and ciliaOTHER (CG500 excluding CCGS genes; **e.**). Curves show the observed high/low groups, shaded areas indicate 95% confidence intervals. Hazard ratios and displayed *p*-values were obtained from multivariable Cox proportional hazards models adjusted for age, sex, pathological stage and CMS subtype; *p*-values were Benjamini–Hochberg-adjusted across the tested signatures. **f.** CCGS GSVA score across pathological tumour stages (Stage I-IV) in the TCGA cohort, coloured by a GSVA-based erdafitinib sensitivity score derived from the erdafitinib-associated differential expression gene set identified in Fig. 1. The reported *p*-value is from a linear model with stage encoded as an ordered numeric variable (I-IV = 1-4). **g-h.** KM plots of PFS stratified by CCGS GSVA score within early-stage (Stage I-II; **g.**) and late-stage (Stage III-IV; **h.**) tumours in the TCGA cohort. Stratification was performed using the same score cut-offs as in **c**. Curves show the observed high/low groups. Hazard ratios and displayed *p*-values were obtained from multivariable Cox proportional hazards models adjusted for age, sex and CMS subtype. **i.** Distribution of CCGS GSVA scores across Consensus Molecular Subtypes (CMS1-4) in the TCGA cohort. Statistical significance was assessed using Dunn’s post-hoc test with BH correction (adjusted *p*-value shown). **j.** Gene Ontology (GO) Biological Process enrichment of genes differentially expressed between tumours in the top and bottom quartiles (25%) of CCGS GSVA score in TCGA. Dot size indicates gene count and colour reflects *−* log_10_(adjusted *p*-value). Selected pathways are shown. **k.** Volcano plot of the same differential expression analysis. Log_2_ fold change (high versus low) is shown on the x-axis and *−* log_10_(BH-adjusted *p*-value) on the y-axis. Wnt-associated genes and Wnt-related transcription factors are highlighted. **l.** CCGS GSVA scores in TCGA colorectal cancer stratified by *APC* alteration status. Each point represents one sample; horizontal bars indicate the median. Associated *p*-value using a two-sided Wilcoxon rank-sum test is shown. **m.** Volcano plot as in **k**, with FGFR signalling genes and erdafitinib targets highlighted.

### The ciliogenic cell state maps to Wnt-associated tumour remodelling in CRC

In the context of the Consensus Molecular Subtype (CMS) framework [22], a widely used transcriptomic classification system in CRC, CCGS was most strongly represented in CMS4, consistent with its association with advanced and aggressive disease states (Figure 2i; Supplementary Figure 10b,d). Interestingly, it was also enriched in CMS2, the canonical Wnt-active subtype. This pattern suggested that the identified cell state might not only mark progression, but also relate to Wnt-associated tumour cell reprogramming. To test this hypothesis, we compared CCGS-high and -low tumours and found that high-scoring samples were enriched for Wnt signalling, including LEF1 expression, extracellular matrix organisation, developmental pathways, and stem cell differentiation, while showing reduced expression of programmes linked to gastrointestinal epithelial maintenance (Figure 2j-k; Supplementary Figure 11a). Consistently, the signature was further associated with alteration of the major upstream Wnt regulator APC, both in TCGA and in our discovery cohort (Figure 2l; Supplementary Figure 10c). Although erdafitinib is approved for FGFR3-altered urothelial carcinoma [23], we observed increased FGFR1 expression in CCGS-high tumours (Figure 2m). Rather than reflecting canonical ciliogenesis, the observed transcriptional programme is consistent with selective activation of cilia-associated genes within a broader Wntand FGFR-linked tumour state, in line with reports that these pathways can induce context-specific ciliary gene module expression [24, 25]. Together, these findings suggest that the ciliogenic programme occurs within a broader Wntand FGFR-driven tumour state marked by extracellular remodelling, developmental signalling, and reduced gastrointestinal epithelial identity.

### Single-cell analysis places the ciliogenic cell state along CRC progression toward non-canonical tumour states

Considering this developmental link, we next investigated the ciliogenic signature in the context of CRC tumour progression at single-cell resolution. Using a recently published scRNA-seq dataset [8] comprising matched normal tissue, primary tumours and metastases, we found increased CCGS expression from normal to primary to metastatic disease for most patients (Figure 3a-c, Supplementary Figure 12a-d). After confirming CCGS expression in tumour cells, we subsequently explored a CCGS-high versus -low comparison at single-cell resolution. CCGS-high malignant cells again showed enrichment of Wnt-related programmes, including LEF1 expression, together with increased expression of FGFR1 and FGFR2, two direct targets of erdafitinib (Figure 3d-f, Supplementary Figure 13a-c).

**Figure 3:**
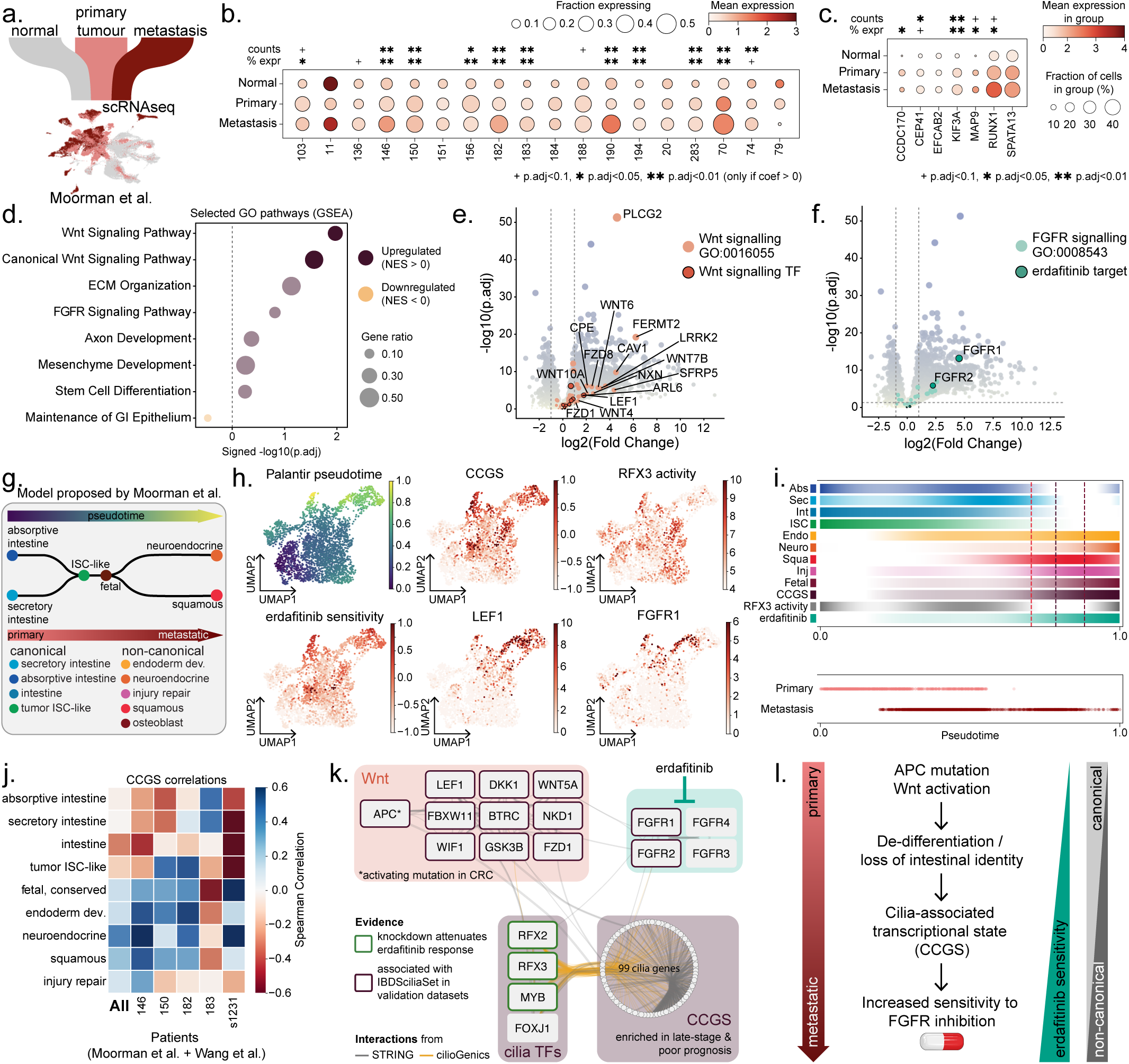
Single-cell analysis places the ciliogenic cell state along colorectal cancer progression toward non-canonical tumour states. **a.** scRNAseq dataset from Moorman et al. consisting of matched normal, primary and metastatic tissue across 31 patients [8]. **b.** Dot plot showing mean CCGS expression across patients stratified by sample type (n = 84 of 99 genes measured in Moorman et al.). Dot size indicates the percentage of cells expressing CCGS genes, and colour indicates mean expression. Stars denote Benjamini-Hochberg-adjusted *p*-values from patient-specific linear models testing for increasing CCGS expression or percentage of CCGS-expressing cells across progression states. Only patients with all sample types represented were included (n = 18 patients). **c.** Dot plot showing mean expression of the highest expressed CCGS genes across patients stratified by sample type. Dot size indicates the percentage of cells expressing each gene, and colour indicates mean expression. Stars denote Benjamini-Hochberg-adjusted *p*-values from gene-specific linear models testing for increasing mean expression or percentage of expressing cells across progression states. Only patients with all sample types represented were included (n = 18 patients). **d.** Gene set enrichment analysis (GSEA) of selected GO Biological Process pathways from differential expression between the top and bottom 20% of CCGS-expressing pseudobulk samples. The x-axis shows *−* log_10_(Benjamini– Hochberg-adjusted *p*-values). Dot size indicates gene ratio, and fill colour indicates directionality, with lower opacity marking pathways below the significance threshold. **e–f.** Volcano plot of differential expression analysis between top and bottom 20% of CCGS-expressing pseudobulk samples (n = 18 matched samples). Log_2_ fold change (high versus low) is shown on the x-axis and *−* log_10_(Benjamini-Hochberg-adjusted *p*value) on the y-axis. **e.** Wnt-associated genes and Wnt-related transcription factors are highlighted. **f.** FGFR signalling genes and erdafitinib targets are highlighted. **g.** Tumour progression model as proposed by Moorman et al. [8] **h.** UMAPs of cells from patient KG146 (n = 3351 cells). Cells are coloured by Palantir pseudotime (top left), Hotspot module score of CCGS (top middle), decoupleR-inferred RFX3 transcription factor activity (top right), Hotspot module score of predicted erdafitinib sensitivity (bottom left, GSVA-derived transcriptomic score based on the erdafitinib-associated differential expression gene set from Figure 1), LEF1 expression (bottom middle), and FGFR1 expression (bottom right). **i.** Gene module score trends across pseudotime in all tumour cells from patient KG146 (top). Each row represents the module score, or RFX3 transcription factor activity, scaled from the 15th percentile (white) to the maximum value (highest saturation). Dashed lines indicate the 75th percentile of each score. Dots indicate tumour cell type along palantir pseudotime (bottom). **j.** Correlations of CCGS module score with the cell states defined by Moorman et al. across all patients, in individual patients KG146, KG150, KG182, and KG183, and in patient s1321 from Wang et al.[35]. **k.** Network visualisation of reported interactions from STRING[30] and cilioGenics[21] among selected Wnt-related genes, cilia-associated transcription factors (TFs), CCGS genes and erdafitinib targets. Elements supported by evidence in this study are indicated. **l.** Proposed model of a progression-associated CRC state linked to increased sensitivity to erdafitinib.

In the proposed model of CRC progression associated with this dataset [8], tumour cells move along pseudotime from canonical intestinal states through a fetal progenitor intermediate towards non-canonical states (Figure 3g). Within this context, we found that the ciliogenic cell state increased along pseudotime with the emergence of noncanonical states and coincided with predicted erdafitinib sensitivity, further placing the cell state along a progression-associated tumour-cell trajectory (Figure 3h-i; Supplementary Figure 13d). CCGS expression correlated preferentially with fetal and noncanonical states, but not with canonical intestinal states (Figure 3j). RFX3 activity rose earlier along the same trajectory and was associated with the earlier ISC-like state, consistent with an upstream regulatory role in this programme (Figure 3h-i; Supplementary Figure 13d-e). Together, these results place the ciliogenic programme within a transition away from canonical intestinal identity towards fetal and non-canonical tumour states.

## 3 Discussion

Collectively, these data suggest a model in which APC/Wnt-associated developmental remodelling during colorectal tumorigenesis [26, 27, 28] facilitates downstream acquisition of a ciliogenic cell state including motile cilia gene expression atypical of normal intestinal epithelial cell states [24, 18, 21]. This is analogous to Wnt-driven ciliogenesis in developmental contexts [24, 25, 29], and coincides with ex vivo sensitivity to the FGFR-inhibitor erdafitinib (Figure 3l). Consistent with this interpretation, network analysis positions Wnt-related genes, cilia-associated transcription factors, CCGS genes, and FGFR family members within a shared interaction landscape (Figure 3k) [30, 21]. Erdafitinib shows tumour-agnostic activity in tumours harbouring FGFR fusions or activating mutations (RAGNAR [31]; NCI-MATCH K2 [32]; FIGHT-207 [33]), but such alterations are rare in colorectal cancer, and genetic FGFR amplification, the more common type of FGFR alteration in CRC, has not emerged as a robust predictor of FGFR-inhibitor benefit (NCI-MATCH K1 [34]). Complementary to this genomic axis, we identify CRC patient stratification by ex vivo functional sensitivity to erdafitinib, and report a transcriptional ciliogenic signature (CCGS) that reflects the underlying FGFR-inhibitor-sensitive cancer cell state.

Pending prospective validation, our findings nominate a patient-stratified repurposing opportunity for an approved FGFR inhibitor in a defined colorectal cancer population, and more broadly illustrate how functionally guided, multi-omic profiling can surface actionable tumour cell states beyond the reach of genomic classification.

## 5 Methods

### 5.1 Patient cohort

This study was conducted as a prospective, non-randomised observational clinical study. The primary outcome was a feasibility assessment of parallel ex vivo drug screening and transcriptomics in patients with CRC. The study was conducted in compliance with the Swiss law, and Swiss regulatory authority requirements, as applicable. Ethical approval: **BASEC ID 2019-01441**. Informed consent was obtained from all participants and/or their legal guardians. Patients with CRC undergoing routine diagnostic or therapeutic procedures at University Hospital Zurich (USZ), Stadtspital Waid und Triemli (STZ), Spital Männedorf (MAE), Klinik Hirslanden (HRZ), Kantonsspital Baselland (KSBL), or Claraspital Basel (STC) were eligible for the study. A detailed list of patients included in the final cohort is provided in Supplementary Table S1. Clinical annotations and follow-up data were collected as part of clinical routine during the same time period. Several clinical parameters, including sex, mutation status, and MSI status were supplemented with RNA-derived annotations (see Section 5.4).

### 5.2 Tissue dissociation

Patient tissue was manually dissected using a scalpel, then enzymatically dissociated in 50% Accutase (STEMCELL Technologies), collagenase IV (1 mg/mL, SigmaAldrich), hyaluronidase (500 ng/mL, Sigma-Aldrich), and DNase I (0.1 *µ*g/mL, Axon Lab AG) in RPMI 1640 + GlutaMAX cell culture medium (Gibco) for 1 hour on a rotator (MACS Mixer) at 37*^◦^*C and 5% CO_2_. The resulting cell suspension was passed through a 70 *µ*m cell strainer and washed with medium. Red blood cells were lysed using RBC lysis buffer (BioLegend) for 10 minutes at room temperature before the cells were resuspended in RPMI 1640 + GlutaMAX medium supplemented with 10% fetal bovine serum (FBS, Gibco) and Antibiotic-Antimycotic (1X, Gibco).

Peripheral blood was diluted 1:1 with phosphate-buffered saline (PBS, Gibco). Peripheral blood mononuclear cells (PBMCs) were isolated using Lymphoprep (STEMCELL Technologies) by density centrifugation at 800 *×* g for 30 minutes, following the manufacturer’s instructions. Red blood cells were lysed using RBC lysis buffer (BioLegend) for 10 minutes at room temperature before PBMCs were resuspended in RPMI 1640 + GlutaMAX medium supplemented with 10% FBS. Cells were counted using the Countess II Cell Counter (Thermo Fisher) according to the manufacturer’s protocol.

### 5.3 Image-based ex vivo drug screening

#### 5.3.1 Drug screening library preparation and storage

The drug library was selected in consultation with CRC-treating physicians to include US Food and Drug Administration (FDA)-approved compounds for CRC and other solid tumours. The complete list of compounds used in this study can be found in Supplementary Table S2. Drugs were prepared as 5 mM stock solutions and dispensed into CellCarrier 384 Ultra microplates (PerkinElmer, 6057300) using an Echo 550 liquid handler (Labcyte) in a randomised layout to minimise plate effects. Unless otherwise stated, all small molecules were tested at a final concentration of 10 *µ*M and antibodies at 10 *µ*g/mL, in 4 replicates per drug or drug combination, and 16 replicates for the DMSO control.

#### 5.3.2 Cell seeding and drug testing

Freshly dissociated cells were mixed with patient-matched PBMCs at a 1:1 ratio and then seeded into prepared drug plates in 50 *µ*L at 0.5*×*10^4^ to 1*×*10^4^ cells per well. Cells were incubated with drugs for 24 hours at 37*^◦^*C, 5% CO_2_. The medium was then aspirated using a microplate washer (Tecan), and cells were fixed for 10 minutes at room temperature with 20 *µ*L/well of a periodate–lysine–formaldehyde fixative (75 mM lysine, Sigma Aldrich L5626-100G; 2.5 mg/mL sodium periodate, Sigma Aldrich 30323-100G; and 1.25% (v/v) formalin, 37% formaldehyde solution, Sigma Aldrich F8775-500ML in PBS). After fixation, the fixative was removed and 70 *µ*L of PBS/well was added. Plates were then stored at 4*^◦^*C for up to two weeks before staining.

#### 5.3.3 Immunofluorescence staining

For staining, PBS was aspirated and cells were blocked, permeabilised, and stained for DNA using 20 *µ*L of PBS containing 5% FBS (Gibco/ThermoFisher 10270106), 2 *µ*g/mL 4’,6-diamidino-2-phenylindole (DAPI, BioLegend 422801), and 0.1% (v/v) Triton-X100 (Sigma Aldrich T8787) for 30 minutes at room temperature. The blocking solution was then aspirated, and cells were stained with fluorescently labelled antibodies diluted in PBS with 10 mg/mL bovine serum albumin (BSA, Sigma Aldrich A7906) for 2 hours at room temperature or overnight at 4*^◦^*C. For details on the antibodies used and staining panel per sample, see Supplementary Tables S2. Finally, the antibody-containing solution was aspirated and PBS was added to the cells. The plate was stored at 4*^◦^*C until imaging.

#### 5.3.4 Confocal imaging and image analysis

All samples were imaged using an automated spinning-disk confocal microscope (PerkinElmer Operetta CLS or PerkinElmer Opera Phenix, see Supplementary Table S2) at 20*×* magnification, acquiring 25 images per well to cover the entire well area. Selected images were acquired at 40*×* magnification for visualisation. Imaging was performed using five channels with non-overlapping excitation/emission filters to capture distinct cellular features. Brightfield imaging (transmission/650–760 nm) was used in Channel 1 to visualise general cell shape and texture. Channel 2 (405 nm/435– 480 nm) was used for nuclear staining with DAPI. Cancer markers were imaged in Channel 3 (488 nm/500–550 nm) and Channel 4 (561 nm/570–630 nm). Immune cells (CD45) were captured in Channel 5 (640 nm/650–760 nm). Single-cell segmentation was performed using the DAPI channel in CellProfiler 2.2.0. Pipeline parameters were adjusted for each sample to accommodate variations in nuclear intensity, size, and the presence of cell clusters. Nuclear expansion was applied to evaluate cytoplasmic features, including marker expression. Subsequent image analysis was conducted in MATLAB R2019a–R2023a.

#### 5.3.5 Cell classification using deep learning

Segmented cells were classified using convolutional neural networks (CNNs) based on a ResNet18 architecture (Supplementary Figure 3f), trained as previously described [36, 37, 38]. The first model, CNN1, was a two-class classifier trained on the DAPI and brightfield channels to distinguish cells from debris. Training was performed on 68,444 manually curated, class-balanced 48*×*48 pixel single-cell crops from all available samples, with 15% of the dataset used for validation. Accuracy and persample accuracy were evaluated on an independent, manually curated dataset of 20 single-cell crops per class per sample (Supplementary Figure 3b-c). Objects classified as debris with a probability *>* 85% were excluded from further analysis.

Next, CNN2, a three-class classifier, was trained to categorise cells as ’malignant’, ’leukocyte’, or ’other’ using all five imaging channels. This model was trained on 76,443 manually curated, class-balanced 150*×*150 pixel single-cell crops, with marker expression guiding manual curation. Again, 15% of the dataset was used for validation, and accuracy and per-sample accuracy were verified on an independent set of 20 singlecell crops per class per sample (Supplementary Figure 3d-e). Final classification was assigned based on the highest predicted class probability.

Cell composition per sample was calculated as the mean fraction of each cell type in DMSO control wells. Samples with a malignant fraction *<* 0.5% were excluded from further analysis.

#### 5.3.6 Calculation of drug response

Two different cancer-specific drug response readouts, RCN and RCF, were calculated as follows:

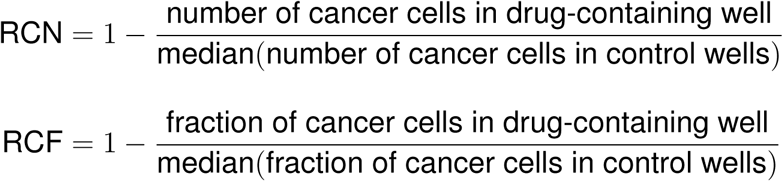

In both cases, a positive score reflects a desirable outcome, namely a reduction in tumour cells. However, RCN focuses solely on tumour cell counts and assigns positive values to any cytotoxic treatment, regardless of its specificity to malignant cells. In contrast, RCF only yields a positive score when the reduction in malignant cells exceeds that of non-malignant cells, thereby highlighting compounds that selectively target malignant cells (Supplementary Figure 5a). For that reason, RCF was used in further analyses. Statistical significance of ex vivo response was evaluated using a two-sided two-sample *t*-test, comparing each drug to its respective control (DMSO for small molecules and isotype control for antibodies), with multiple testing correction applied using the Benjamini–Hochberg (BH) method.

For further analyses, phenotypes were considered a ’response’ (R) with RCF *>* 0 and BH-adjusted *p*-value *<* 0.05 across drugs, all other phenotypes were classed as ’no response’ (NR).

For visualisation purposes in Figure 1a and Supplementary Figure 4, LASSO regression was used to infer missing drug responses that were not experimentally measured. Drugs with partial missingness were modelled using fully observed drugs as predictors, restricting training to patients with observed responses (minimum n = 30). Five-fold cross-validation was used, and imputation was performed only if the Pearson correlation between cross-validated predicted and observed responses exceeded 0.6. Missing values were replaced with model predictions constrained to the empirical range of observed responses. Imputed values were used exclusively for visualisation and not for statistical analyses or response classification.

### 5.4 Transcriptomics

#### 5.4.1 RNA extraction

A pellet containing approximately 1 million cells was collected after tissue dissociation, resuspended in 350 *µ*L of TRIzol reagent (*Ambion, ThermoFisher* 15596026), and stored at *−*80*^◦^*C until RNA extraction. RNA was extracted using the Zymo Directzol RNA Miniprep Kit (*Zymo* R2052), following the manufacturer’s instructions. RNA concentration and integrity were assessed using a TapeStation with RNA reagents (*Agilent Technologies*).

#### 5.4.2 Library preparation and sequencing

Libraries were prepared using the Illumina TruSeq Stranded Total RNA kit (with ribosomal depletion), and sequencing was performed on an Illumina NovaSeq 6000 platform using single-end, 100 bp reads. The sequencing depth was approximately 40 million reads per sample. Raw sequencing data were obtained in FastQ format.

#### 5.4.3 Processing of raw RNA-seq reads

Raw reads were aligned to the human reference genome GRCh38.p13 (release 42) using STAR (version 2.4.2b), and gene expression was quantified using featureCounts [39]. Read quality was assessed using FastQC (version 0.11.9), and low-quality samples or outliers were excluded from further analysis.

#### 5.4.4 Mutational profiling using transcriptomics

The mutation-calling pipeline followed the Genome Analysis Toolkit (GATK) best practices [40]. The workflow and associated quality control are illustrated in Supplementary Figure 14 and consisted of three main steps: RNA-seq preprocessing, mutation calling, and variant effect prediction.

##### RNA-seq preprocessing

Reads were aligned to the GRCh38.p14 human reference genome (*Ensembl*, version 111) using STAR (version 2.5.2b). The resulting alignments were processed with Picard (version 3.1.1) using the MarkDuplicates tool to identify PCR duplicates. GATK (version 4.5.0) was used for exon segmentation (SplitNCigarReads) and base quality score recalibration (BQSR), performed using BaseRecalibrator and ApplyBQSR. Known SNPs were excluded from recalibration using NCBI dbSNP (build 156, GRCh38.p14).

##### Mutation calling

Somatic short mutations were identified using the MuTect2 variant caller in tumour-only mode, due to the absence of matched normal samples. A panel of normals and a germline resource from GATK GRCh38 were included to improve specificity. Additional filtering was applied using FilterMutectCalls to remove technical artefacts, nonsomatic mutations, and sequencing errors. Only variants previously documented in the literature were retained. Because variant detection was performed from RNA-seq in tumour-only mode, the resulting calls should be interpreted as transcript-supported variant calls rather than comprehensive genomic mutation profiles. This approach has known limitations, including dependence on gene expression and transcript coverage, potential allelic imbalance, and reduced sensitivity for truncating alterations that may be depleted at the RNA level [41, 42]; accordingly, mutation frequencies for some genes may be underestimated.

##### Variant effect prediction

The impact of somatic mutations was assessed using the Variant Effect Predictor (VEP, Ensembl version 111). Only mutations classified as high or moderate impact were retained, with a focus on genes commonly altered in CRC, as documented in the COSMIC large intestine mutation census [43]. Further analysis included only genes mutated in at least three patients.

#### 5.4.5 Quality control and filtering

The total unfiltered count of genes detected per sample is shown in Supplementary Figure 6a. For downstream analysis, only protein-coding and long non-coding RNAs were retained. Genes were filtered to include those with a minimum read count of 10 in at least 15 samples, resulting in 18,497 unique genes used for analysis.

#### 5.4.6 Deconvolution, MSI status, and CMS prediction

Raw counts were variance-stabilised using the vst function from DESeq2 (version 1.38.3) and batch-corrected using removeBatchEffect from limma (version 3.54.2) prior to MSI and CMS prediction, deconvolution, and sex derivation.

Bulk RNA-seq deconvolution was performed using DigitalDLSorteR (version 0.3.1) with the colorectal-specific pre-trained model colorectal.li from DigitalDLSorteRmodels (version 0.1.0). The deconvolution results were validated using nine samples with immunofluorescence-derived, tumour-only (PBMC-free) fractions (Supplementary Figure 6b-c).

MSI status was predicted using the PreMSIm package (version 1.0) [44], and CMS subtypes were assigned using CMScaller (version 2.0.1) [45]. Patient sex was inferred using Y-chromosome-specific *RPS4Y1* expression as previously described [46].

#### 5.4.7 Gene set variation analysis

Raw count data were variance-stabilisation transformed (VST) using the vst function from DESeq2 (version 1.38.3). Batch effects and CRC fraction (as determined by DigitalDLSorteR, see above) were regressed out using the removeBatchEffect function from limma. Gene set variation analysis (GSVA) was performed using the gsva function from GSVA (version 1.53.8) with the default Gaussian kernel (kcdf = ”Gaussian”) for VST-transformed data. Gene sets were supplied with a minimum size of five genes per set.

### 5.5 Integrative analysis of drug response and gene expression

Only drugs with more than 20 non-missing response measurements and at least five responding samples (RCF *>* 0, BH-adjusted *p*-value *<* 0.05) were retained for integrative analyses. Drug response was binarised per sample as responder (R) or nonresponder (NR), as described above. For each drug, only samples with non-missing response were included in downstream analyses.

#### 5.5.1 Differential expression analysis (DEA)

Differential expression analysis was performed separately for each drug using DESeq2 (version 1.38.3) on raw count data. Only protein-coding genes were retained. Batch and CRC fraction (determined by deconvolution, as described above) were included as covariates. The DESeq function was run with minReplicatesForReplace = 5. Log_2_ fold changes (FC) were shrunk using lfcShrink with the apeglm method.

#### 5.5.2 Gene-drug network and Louvain communities

Differential expression results across all drugs were integrated into a bipartite gene-drug association network. For each drug, genes were retained if they were upregulated in responders (shrunken log_2_ FC *>*1 and BH-adjusted *p*-value *<*0.05); to limit the influence of very large gene sets, edges were further restricted to the top 300 genes per drug ranked by log_2_ FC. The network was constructed as an undirected graph with drugs and genes as nodes and drug-gene associations as edges, and edge weights were defined as *|* log_2_ FC*|*. Communities were identified using Louvain modularity optimisation as implemented in igraph (cluster louvain), and the resulting community membership was assigned to all nodes (drugs and genes).

#### 5.5.3 GO-term enrichment

Gene Ontology enrichment analysis was performed across Louvain communities using clusterProfiler (version 4.6.2). Enrichment was computed with enrichGO via compareCluster, testing all ontologies (BP, CC and MF) unless otherwise specified. The background universe was defined as all detected protein-coding genes included in the differential expression analyses. BH-adjusted *p*-values shown.

#### 5.5.4 Transcription factor activity inference

Transcription factor activity was inferred using the decoupleR package (version 2.4.0) [47]. The transcription factor gene network from CollecTRI, as previously described [48], and supplemented with a cilia-specific transcription factor annotation, as described in *CilioGenics* [21], was used. The run ulm function was used to infer transcription factor activity based on shrunken log_2_ FC from DEA analysis, resulting in an activation score (the *t*-value of the univariate linear model slope).

#### 5.5.5 Definition of the CCGS

To derive a compact cilia-associated module aligned with the drug-responder signal, we define a geneset using the IBDS bulk RNA-seq cohort (n = 56). Using variancestabilised, batchand tumour-fraction–corrected expression values, a gene–gene Spearman correlation matrix for CG500 genes was performed and then hierarchically clustered (Pearson correlation distance, Ward’s linkage).

Cluster resolution was assessed across *k* = 15–30 using mean silhouette width, and *k* = 21 was selected as a local optimum yielding interpretable module sizes. The cluster containing the largest number of Louvain community 5 cilia-associated genes was defined as the CCGS (n = 99 genes) and used for downstream analyses.

### 5.6 siRNA knockdowns in GP2d

#### 5.6.1 Cell culture

The GP2d human colon adenocarcinoma cell line (Sigma Aldrich, acc no 95090714, lot 23F031, P57, directly purchased from the vendor) was cultured in standard culture medium (DMEM/F12 + GlutaMAX, 10% FBS, 1% penicillin–streptomycin, all from Gibco) at 37*^◦^*C and 5% CO_2_. Cells were passaged using TrypLE Express Enzyme (Gibco).

#### 5.6.2 siRNA knockdown

All siRNAs used in this study were part of the MISSION esiRNA library (SIGMA, Supplementary Table S3). FLUC esiRNA (EHUFLUC), targeting firefly luciferase, was used as a negative control, and KIF11 esiRNA (EHU019931) was used as a positive control for transfection and viability. siRNAs were transfected at 10 ng per well in 384-well plates (used for imaging and drug incubation) and 40 ng per well in 96well plates (used for RNA extraction and quantitative real-time PCR (qRT–PCR)) with Lipofectamine RNAiMAX (Invitrogen, 13778075) in Opti-MEM (Gibco, 31985062). For 384-well plates, siRNAs were dispensed using a Labcyte Echo liquid handler in a randomised layout to control for plate effects and stored at *−*20*^◦^*C until use. Cells were incubated for 48 hours at 37*^◦^*C and 5% CO_2_ after transfection, prior to RNA extraction or drug treatment. For drug treatment, drugs were added on top using an INTEGRA MINI-96 liquid handler and incubated for a further 24 hours before fixation and staining (as described in Section 5.3). For RNA extraction, cells were washed and 150 *µ*L of TRIzol LS (Thermo Fisher, 10296028) was added per well in a 96-well plate, which was then stored at *−*20*^◦^*C until extraction.

#### 5.6.3 Incucyte live-cell imaging

GP2d cells were reverse-transfected as described above in a 96-well Ultra microplate (PerkinElmer), with 20,000 cells seeded per well. Real-time confluence of cell cultures (n = 3 replicate wells per condition) was monitored by imaging every 2 hours for 72 hours at 10*×* magnification in the phase-contrast channel using the Incucyte S3 live-cell analysis system (Sartorius). Automatic image segmentation and confluence analysis were performed using the Incucyte Base Analysis Software (version 2020B). Relative confluence was calculated by normalising values to the first time point (2 hours).

#### 5.6.4 siRNA knockdown efficiency by quantitative real-time PCR

Knockdown efficiency was measured using TaqMan Gene Expression Assays (Thermo Fisher, see Supplementary Table S3) with TaqMan Fast Advanced Master Mix, no UNG (Thermo Fisher, A44360) on a QuantStudio 3 Real-Time PCR System (Applied Biosystems, A28567). Total RNA was extracted using the Direct-zol RNA MicroPrep Kit (Zymo Research, R2062) and quantified with a Qubit 4 fluorometer (Thermo Fisher). cDNA was synthesised using the iScript cDNA Synthesis Kit (Bio-Rad, 1708890). For each TaqMan assay (n = 4 biological replicates), 40 ng of cDNA per sample were used. Knockdown efficiency was calculated based on Δ*C_t_* values, subtracting the *C_t_* of the endogenous control (18S rRNA). ΔΔ*C_t_* values were obtained by subtracting the Δ*C_t_* of the FLUC control from that of the target siRNA. Percent knockdown was calculated as previously described [49]:

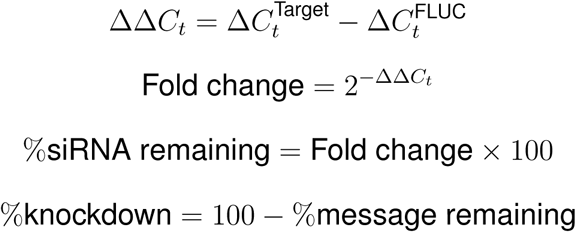

#### 5.6.5 Calculating the siRNA knockdown drug responses

siRNA- and drug-treated GP2d cells were stained with DAPI and cleaved Caspase 3 (see Supplementary Table S2). All drug response calculations were based on the viable total cell number, defined as clCASP3*^−^* cells per well. Each drug condition was tested in four replicates. siRNAs were excluded from analysis if they were toxic (relative confluence *<* 3 after 72 hours) or if knockdown efficiency was inconclusive, insignificant, or undetectable by qPCR.

For each siRNA-drug condition, drug response was calculated as the ratio of viable cell number to the median viable cell number in matched DMSO control wells. To quantify siRNA-specific effects on drug sensitivity, values were normalised to the FLUC control by computing (FLUC-normalised response / siRNA response). Values *<* 1 indicated attenuated drug response, while values *>* 1 indicated increased drug sensitivity.

### 5.7 Publicly available datasets

#### 5.7.1 CilioGenics

For improved cilia-associated gene annotation, a gene list with associated scores, as described by Pir et al., was downloaded from the CilioGenics website (https://ciliogenics.com/, accessed on 8 November 2024) [21]. For the purposes of our analysis, the top 500 cilia-associated genes (ranked by CilioGenics score) were assumed cilia-related and referred to as CG500.

#### 5.7.2 The Cancer Genome Atlas (TCGA)

The TCGA colorectal cancer cohort (COADREAD Pan-Cancer Atlas, n = 592) was accessed using the TCGAretriever package (version 1.9.1). Log-normalised expression values were generated using scuttle::logNormCounts. Tumour fraction was estimated using DigitalDLSorteR and regressed out from log-normalised expression values using limma::removeBatchEffect, resulting in the matrix used for downstream analyses.

Gene set variation analysis (GSVA) was performed using the GSVA package with a Gaussian kernel (kcdf = ”Gaussian”) and a minimum gene set size of five genes. Gene sets included CCGS, CG500, and ciliaOTHER (CG500 excluding CCGS genes). For the prediction of erdafitinib sensitivity, we used all differentially expressed (shrunken log_2_ FC *>*1 and BH-adjusted *p*-value *<*0.05) genes associated with erdafitinib responders in IBDS CRC, as used for Figure 1.

##### Survival analysis

Associations between GSVA scores and progression-free survival (PFS) were assessed in the TCGA colorectal cancer cohort using Cox proportional hazards models. Optimal cut-offs for dichotomising GSVA scores were determined in the full cohort using the maximally selected rank statistic implemented in surv cutpoint (minprop = 0.2), and the resulting thresholds were applied to all subsequent subgroup analyses. Samples were classified into high and low groups for each signature. Primary models included the dichotomised signature together with age, sex, pathological stage and CMS subtype. Stage-stratified models included age, sex and CMS subtype. Hazard ratios and *p*-values for the signature term were extracted from model summaries using the Wald test. For the primary analysis across tested signatures, *p*-values were adjusted using the Benjamini-Hochberg method. Kaplan-Meier curves were generated for the observed high and low groups using survfit and visualised with ggsurvplot.

##### Differential expression and GO enrichment

To characterise transcriptional programmes associated with high CCGS activity in TCGA, samples were stratified by CCGS GSVA score into high (top quartile, *≥* 75th percentile) and low (bottom quartile, *≤* 25th percentile) groups; samples in the interquartile range were excluded. Differential expression was performed using limma on log-normalised expression values, modelling CCGS group and tumour fraction as covariates.

Linear models were fitted with lmFit and empirical Bayes moderation was applied with eBayes. Significance was assessed using BH-adjusted *p*-values.

For Gene Ontology (GO) enrichment, genes upregulated in high-scoring tumours (BH-adjusted *p <* 0.01 and log_2_ FC *>* 1) and downregulated in high-scoring tumours (BH-adjusted *p <* 0.01 and log_2_ FC *< −*1) were analysed separately using clusterProfiler::compareCluster with enrichGO (Biological Process ontology; pvalueCutoff = 0.01; OrgDb = org.Hs.eg.db; keyType = ENTREZID; readable = TRUE). For volcano plot annotation, genes belonging to curated pathways (e.g. Wnt signalling and FGFR signalling) were retrieved from Gene Ontology annotations using org.Hs.eg.db. Transcription factors within the Wnt-associated gene set were defined based on overlap with the CollecTRI transcription factor–target network [48]. Selected drug target genes were obtained from the STITCH database [50, 51] and highlighted accordingly.

#### 5.7.3 Single-cell RNA sequencing: Broguire et al

A publicly available single-cell RNA-seq dataset from 26 CRC patients was downloaded from https://crc-tme.com/Downloads/OriginalTumorsAndMonocultures_ scRNAseq.zip [19], including Scanorama embedding. Counts were log-normalised for analysis.

#### 5.7.4 Single-cell RNA sequencing: Moorman et al

Single-cell RNA-seq data from 31 patients with matched normal colon, primary, and metastatic CRC were downloaded from https://dp-lab-data-public.s3. us-east-1.amazonaws.com/progressive-plasticity-crc-metastasis/data. tar.gz [8]. Analyses focused on the datasets containing all epithelial cells (AllEpithelial.h5ad file), epithelial tumour cells of the entire cohort (Tumor.h5ad file), and epithelial tumour cells of individual patients.

##### Association of CCGS expression in matched normal, primary, and metastatic samples

To assess changes in CCGS expression across disease progression (normal, primary tumour, and metastasis), single-cell data were aggregated (mean counts) into patient- and stage-specific pseudobulk profiles. This resulted in 54 samples derived from 18 patients with matched representation across all three stages. Progression-associated expression changes were evaluated using linear modelling approaches at both the gene and patient level. At the gene level, expression of individual CCGS genes was modelled as a function of progression stage within each patient, thereby accounting for patient-specific effects. At the patient level, the mean expression of the CCGS was modelled as a function of progression stage, discretely encoded in order from normal to primary to metastatic, while accounting for gene-specific baseline differences. In addition to mean expression, analyses were performed on the proportion of cells expressing each gene (gene-level) or the gene set (patient-level). Linear models were fitted using statsmodels (version 0.14.6), and resulting *p*-values were adjusted for multiple testing using BH method.

##### Hotspot module score calculation

Hotspot modules (Moorman et al. [8], Supplementary Table S4) were supplemented with gene sets identified in this study (incl. CG500, CCGS and ex vivo erdafitinib sensitivity associated genes from the differential expression analysis in Figure 1 used to calculate predicted erdafitinib sensitivity). Gene module scores were calculated using calculate module scores from Hotspot (version 1.1.3) as described in the original study [8].

##### Pseudobulk differential expression analysis of CCGS-high and CCGS-low tumour cells

All epithelial tumour cells were stratified according to the distribution of the CCGS hotspot module score, with the bottom 20% classified as cilia-low and the top 20% as cilia-high. Pseudobulk profiles were generated by aggregating raw gene expression counts across cells within each sample and cilia group. To ensure robustness, only sample–group combinations comprising at least 30 cells were retained for down-stream analysis (n = 18 samples). Genes with low overall expression were excluded by requiring a minimum of 10 total counts across all pseudobulk samples. Differential expression analysis between CCGS-high and CCGS-low groups was performed using PyDESeq2 (version 0.5.4) with a paired design, including sample identity as a blocking factor to account for inter-sample variability. Log_2_ fold changes were subjected to shrinkage to improve effect size estimation and gene ranking. The resulting ranked gene list was used for gene set enrichment analysis (GSEA) of Gene Ontology biological process (GO:BP) pathways using gseapy (version 1.1.11).

##### Transcription factor activity inference

Transcription factor activity was inferred based on the t-statistic resulting from the epithelial tumour cell pseudobulk differential expression analysis using the transcription factor gene network as described above with the run ulm function from decoupler-py (version 2.1.3).

##### Data visualisation

UMAP embeddings were visualised using scanpy [52] (version 1.11.5) based on previously published UMAP coordinates. For patient KG146 and the dataset from Wang et al. [35], module score trends were calculated as described previously [8]. Briefly, CCGS, erdafitinib sensitivity signature, and RFX3 transcription factor activity module scores were added, and generalised additive models (GAMs) with cubic spline smoothing were fitted as a function of pseudotime. For Wang et al. pseudotime was calculated using diffusion pseudotime [53] as implemented in scanpy. Smoothed scores were calculated across 500 equal-width pseudotime bins and visualised from the 20th percentile to the maximum value. To calculate CCGS and RFX3 activity correlations across cell states identified in Moorman et al., cells were assigned to their respective top-scoring module if they exceeded the 25th percentile of that particular module.

### 5.8 Software

The analysis for ex vivo drug screening, including statistical analysis, was performed using MATLAB (R2020b–R2023a). The processed drug response matrix and associated *p*-values were subsequently imported into R (version 4.2.3) for visualisation and integration with other datasets. All other analyses and visualisations were conducted in R. Analysis of Moorman et al. dataset was done using a CUDA-backed (version 12.9) rapidsai environment (version 25.12) in Python (version 3.13.11).

### 5.9 Statistical analysis

Unless otherwise specified, all statistical tests were two-sided. Ex vivo drug-response analyses used two-sample *t*-tests. Differential expression analyses in DESeq2 used the coefficient-level Wald test. Survival associations were assessed using Cox proportional hazards models, with hazard ratios and *p*-values for the signature term extracted using the Wald test. Pairwise group comparisons used Wilcoxon rank-sum tests where indicated, and multiple-group comparisons used Dunn’s test with BH correction. Linear trends across ordered pathological stages were tested using linear models with stage encoded as an ordered numeric variable. *P* -values were adjusted for multiple testing using the BH method within the relevant family of hypotheses, as specified for each analysis.

## Data availability

All transcriptomics data generated in this study will be deposited in the National Center for Biotechnology Informations Gene Expression Omnibus (GEO; https:// www.ncbi.nlm.nih.gov/geo/). Previously published scRNA-seq datasets analysed in this study were downloaded from http://humantumoratlas.org/publications/hta8_ crc_moorman_2024. The publicly available GRCh38 human reference genome was used to align RNA-seq reads. Protein–protein interaction data were retrieved from STRING (https://string-db.org/). Data provided in supplementary tables include ex vivo drug response of colorectal cancer cells, patient meta-data, details on antibodies used in the study, and gene set definitions. Source data will be provided with this paper. Supplementary information, including ex vivo drug response data, patient metadata, antibody details and gene set definitions, will be made available upon journal publication.

